# DCL1, a Protein that Produces Plant MicroRNA, Coordinates Meristem Activity

**DOI:** 10.1101/001438

**Authors:** Stephen E. Schauer, Teresa A. Golden, Delwin S. Merchant, Biranchi N. Patra, Jean D. Lang, Sumita Ray, Bulbul Chakravarti, Deb Chakravarti, Animesh Ray

**Affiliations:** Institute of Plant Biology, University of Zürich, Zollikerstrasse 107, CH-8008 Zürich Switzerland; Southeastern Oklahoma State University, Department of Biological Sciences, 1405 N. 4^th^ Ave, PMB 4087, Durant, OK 74701; School of Medicine, Case Western Reserve University, 10900 Euclid Avenue, Cleveland, OH 44106; 217 Berkeley St, Rochester, NY 14607; Senomyx, Inc., 4767 Nexus Centre Drive, San Diego, CA 92121

## Abstract

The ubiquity and importance of short duplex RNAs, termed microRNA (miRNA), for normal development in higher eukaryotes are becoming increasingly clear. We had previously shown that reduction-of-function mutations in *Arabidopsis thaliana DCL1* (*DICER-LIKE1*) gene, affecting the nucleus-localized protein that produces 19-25 nucleotides long miRNA species from longer double stranded RNA precursors, cause a delay in flowering by prolonging the period of juvenile organ development. Here we show that *DCL1* transcription is increased at the critical phase of juvenile to reproductive developmental transition, and that DCL1 protein is localized in meristematic cells of the shoot, inflorescence and flowering meristem. DCL1 protein is also expressed in the ovule funiculus, ovule integuments, and in early but not late embryo. Genetic analysis revealed that *DCL1* exerts its effect along the same pathway that involves the floral pathway integrator gene *LEAFY*. Results are most consistent with the idea that DCL1 protein is required in the shoot apical meristem to prevent uncontrolled proliferation of meristematic cells. The expression of DCL1 protein in the early embryo may be either via the transmission of *DCL1* mRNA through the female gametophyte, as suggested from the sporophytic maternal effect of *dcl1-8* on early embryo development, or from *DCL1* mRNA synthesized in early embryo cells off the maternally transmitted allele. The requirement of an active maternally transmitted allele of *DCL1* for normal early embryo development, and the presence of DCL1 protein in the early embryo, together suggest that the synthesis of miRNA in early embryo cells is critical for development, but does not rule out potential maternal contribution of miRNA or its precursor molecules into the embryo.

## Introduction

A form of small regulatory RNA, microRNA (miRNA), controls the expression of target genes at the posttranscriptional level by several mechanisms [1,2]. One mechanism involves the degradation of target mRNA having sequence complementarity to the miRNA. A second mechanism involves inhibition of translation, and requires less stringent sequence complementarity between the miRNA and its target mRNA. Biosynthesis of miRNA is dependent on a class of enzymes known as Dicer. Unlike mammals, which have only one Dicer, plants have multiple Dicer-like enzymes that perform specialized functions. In *Arabidopsis thaliana* there are four *DICER-LIKE* genes, *DCL1*-*DCL4* [3]*. DCL1* is required for the production of most classes of miRNA [4,5]. A mutation (*dcl1-9* allele) that deletes one of two dsRNA binding domains [6] and a P415S substitution allele (*dcl1-7*) that perturbs the RNA helicase domain [7], each of which removes most species of miRNA molecules encoded by Arabidopsis, suggesting that this protein is essential for the processing of miRNA precursors into mature miRNA [4,5,8,9]. While in animals the processing of hairpin-containing long primary precursor transcripts (pri-miRNA) to shorter pre-miRNA requires the participation of another enzyme, Drosha, in plants both steps are catalyzed by the activities of the single enzyme DCL1 [4,10-14]. *DCL1* is neither involved in post-transcriptional silencing of transgenes [9], nor it is necessary for siRNA mediated silencing of viral RNA expression [15-18].

Since their discovery, plant miRNAs were implicated as regulators of developmentally important genes [18]. Examples range from the recognition that developmentally important genes such as *DCL1* and *ARGONAUTE1* (*AGO1*) encode gene products that process miRNA and/or miRNA targets [4,6,7,19-21] to the identification of specific miRNA species with homology to mRNA that encode developmental regulators [2,19,22-24]. AGO1 is the enzyme that cleaves target mRNAs in a complex involving complementary base pairing with its cognate miRNA [25]. The expression of *DCL1* mRNA is itself potentially under feedback negative regulation by DCL1’s action on miR162, a miRNA that contains sequence complementarity to the *DCL1* message [26].

Arabidopsis *DCL1* is preferentially transcribed from the maternally inherited allele, whose activity in the maternal sporophyte and in the embryo is essential for normal embryo development [7, 28]. Activity of *DCL1* is also important for transition from the juvenile to the mature phases of plant development [29]. While complete loss of function mutations of *dcl1* cause embryo lethality [7], reduction of function mutations in *dcl1* produce a range of developmental defects in Arabidopsis [3,6,28-32]. In particular, the *dcl1-9* allele causes unregulated cell division in floral primordia [6], and all weak *dcl1* alleles prolong the period of juvenile development, confound normal ovule morphogenesis, and embryos born on homozygous *dcl1-8* plants exhibit maternal effect pattern formation defects [28,29]. The flowering time delay in homozygous *dcl1-7* or *dcl1-8* mutants is suppressed by a mutation in the *TERMINAL FLOWER1* (*TFL1*) gene [29], which maintains the juvenile phase of the shoot apical meristem (SAM) by negatively regulating *APETALA1* (*AP1*) and *LEAFY* (*LFY*) [33,34]. Both *AP1* and *LFY* encode transcription factors and are classified as floral meristem identity genes [35,36]. Mutations in these two genes are characterized by late flowering, due to the delayed conversion of lateral meristem fate to produce flower, and abnormal floral morphology. The late flowering phenotype of *dcl1-7* or *dcl1-8* is enhanced by *ap1-1*, a partial loss-of-function mutation, suggesting that the targets of *DCL1* might be genes that participate in the conversion of the lateral inflorescence meristem to floral meristem [29].

Here we show that DCL1 protein is localized in the very cell types where *DCL1* has the most phenotypic effects, namely, in cells of the shoot apical meristem (SAM), the inflorescence and floral meristem cells, in ovule integuments and in the early embryo, and we examine the genetic interaction of *dcl1* with *LFY* and other genes important for flower development.

## Results

### DCL1 Protein is Expressed in the Ovule, Early Embryo, and Meristem Cells

We previously showed by *in situ* RNA hybridization analysis that *DCL1* mRNA is present in both inflorescence and floral meristem cells, and that those *dcl1* alleles that delay flowering time also affect this transcript [7]. To test whether *DCL1* transcription is temporally regulated to coincide with transition to flowering, we examined the level of the 6.2 kilo-base (kb) full-length *DCL1* transcript during development by RT-PCR analysis (Fig. 1, A). We detected an increase in the 6.2-kb transcript in RNA samples from wild type plants at a stage immediately before the transition of the lateral meristem to the floral fate. The mRNA level declined upon transition to flowering.

**Figure 1.**
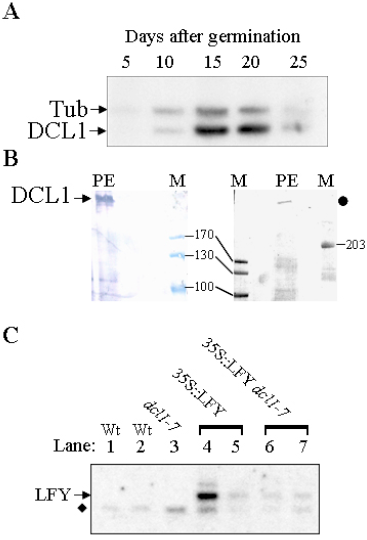
Expression analysis of *DCL1.* A, Temporal expression pattern of the 6.2-kb *DCL1* transcript by RT-PCR with β-1 tubulin (Tub) as an internal control. The ratio of *DCL1* transcript to β-1 tubulin transcript is indicated. B. Western blot analysis of protein extracted from flowers, buds and apices. The blot was probed with affinity purified anti-DCL1 antiserum. Control blots were probed with the preimmune sera, which showed no signal (not shown). Left blot was developed for colorimetric detection, the right blot with chemiluminiscence. Lanes labeled PE contained protein extract, and M had molecular weight markers. The filled circle denotes the position of the high molecular weight band. Numbers are marker sizes (kd). C. RNA gel blot analysis of *LFY* message from wild type (lanes 1, 2), *dcl1-7* (lane 3), *35S::LFY* (lanes 4, 5), and from *35S::LFY dcl1-7* (lanes 6, 7) leaves. Twenty micrograms of total RNA was probed with the full-length cDNA. A second probe used as positive control was cytoplasmic cyclophilin (ROC1) and ethidium bromide stained ribosomal RNA fixed on the membrane post-transfer served as additional loading control (not shown). Arrowhead indicates the position of *LFY* transcript, and the position of the *ROC1* message (loading control) is indicated with a diamond.

To determine whether this pattern of mRNA accumulation in the meristem correlates with expression of DCL1 protein, we raised polyclonal antibody against a protein fragment containing the amino terminal 110 amino acid residues of DCL1 protein. This protein fragment is highly acidic (predicted pI of 3.91) and bears no significant homology to any other known protein from Arabidopsis, including the three other DCL homologs. Western blot analysis with the purified antibody revealed a major band of ∼214 kilodalton (kd) in the soluble fraction of proteins prepared from flowers and buds (Figure 1, B, left), which corresponded to the predicted 214 kd size of DCL1, and a few faint bands of lower molecular weights. The ∼214 kd band was seen consistently in Western blots while the lower molecular weight bands were variable, suggesting the latter bands represent nonspecific degradation products of DCL1. In one western blot, we observed a single strongly reactive band corresponding to ∼430 kd, which either represented a dimeric DCl1 that had failed to denature or a modification product (Figure 1 B, right).

To locate cells that express DCL1 protein, we performed immunocytochemical localization on sections of the SAM in plants undergoing transition to flowering and of lateral meristem and flowers at different developmental stages. DCL1 protein is detected in the shoot apical meristem and emerging leaves of wild type plants (Figure 2 A), and in all three cell layers of inflorescence and floral meristems (Figure 2 E-G).

**Figure 2.**
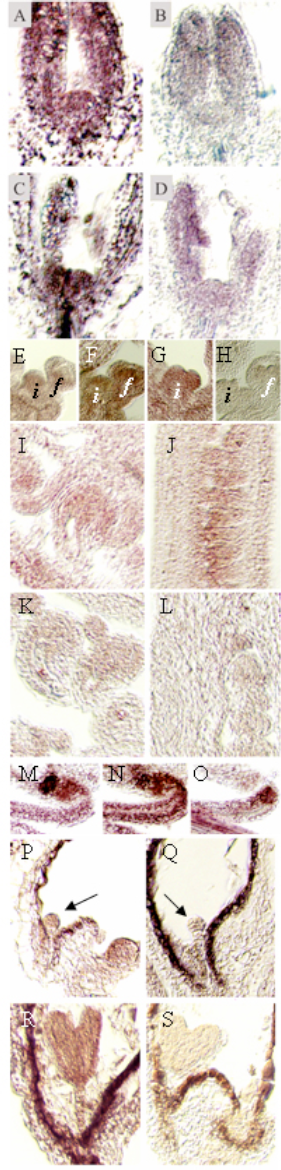
Localization of DCL1 protein. Sections of plant tissues were probed with either DCL1 antiserum (lanes A, C, E-G, I, J, M-O, P, R) or preimmune serum (lanes B, D, H, K, L, Q, S). Abbreviations: i, inflorescence meristem; f, floral meristem. Arrows point to early embryos. Panels A-D are sections of the SAM, E-H of inflorescence and floral meristem, I-O, ovules, and P-S, stages of developing embryos.

DCL1 protein is localized in early ovule primordia (Figure 2 J), in ovule integuments (Figure 2 I, K, L) and in early embryos (Figure 2 P, R). DCL1 protein is detectable in embryos up to the torpedo stage, after which it is undetectable (not shown). These expression patterns of DCL1 protein are consistent with genetic effects of weak *dcl1* mutants that are affected in ovule integument development and strong *dcl1* mutants in early embryo pattern formation, as well as with the patterns of *DCL1* promoter activity described earlier [7,28,30,31].

Strikingly, we were able to localize strong expression of DCL1 protein in the ovule funiculus (Figure 2, M-O), with a punctuate expression pattern in epidermal cells lining the vasculature, which could represent nuclear localization of DCL1 protein. We had previously demonstrated nuclear localization of a green fluorescent protein fused in-frame to an amino terminal DCL1 fragment, and presented evidence for a nuclear function of DCL1 [15]. No morphological defect in the funiculus, however, has yet been seen in *dcl1* mutants.

We investigated the expression of DCL1 protein in transgenic lines harboring *DCL1* cDNA under the control of cauliflower mosaic virus 35S RNA promoter (*35S::DCL1*) described earlier [32]. DCL1 protein is detected in the SAM and emerging leaves of plants expressing the 6.2-kb *DCL1* cDNA (Figure 2, C) at a high level in those lines that show high level expression of DCL1 transcript [32].

### *DCL1* Augments Floral Fate Determination by *LFY*

Results described above together with previous results on the effect of *dcl1* mutation on flowering time [7,29] are consistent with a role for *DCL1* in lateral meristem fate transition. One hypothesis for this role is that negative regulators of reproductive development during juvenile to adult transition are repressed by one or more DCL1-generated miRNA(s). We had previously shown that *dcl1* mutations dramatically enhanced the late flowering effects of weak *ap1-1* allele [29]. Like *AP1*, *LFY* is a major player in determining the floral fate of lateral meristem cells; these two genes are mutually synergistic and *LFY* is a positive regulator of *AP1* transcription. DCL1 could potentially generate a miRNA that down regulates a negative regulator of *LFY*. Alternatively, a specific miRNA could down regulate a negative regulator of a downstream target gene of *LFY*. In this latter role, this miRNA could be a negative regulator of a gene that inhibits flower development. In either case, *dcl1-7*, which removes most species of miRNA, is expected not to affect a null *lfy* phenotype but should enhance a weak *lfy* phenotype. If on the other hand, the miRNA produced by DCL1 affects flowering through a *LFY* independent pathway, then all *dcl1 lfy* double mutants should be synergistically affected in flowering time. To examine these questions, double mutant combinations were made between *dcl1-7* and two alleles of *lfy*.

The strongest allele of *lfy*, *lfy-26*, causes the production of numerous lateral branches, suggesting a conversion of floral meristem into coflorescence shoots, usually with a subtending cauline leaf. Many of these lateral coflorescence branches fail to extend (Fig. 3A). This is in contrast to the wild type where the progression of lateral organs in reproductive development entails two to four elongated coflorescences (subtended by a cauline leaf) followed by flowers (not subtended by cauline leaves; Fig. 3B). As development of the Lfy^-^ primary meristem proceeds, these coflorescence-like ‘flowers’ lose some of their shoot characteristics, including loss of the subtending cauline leaf. Some nearly morphologically normal yet infertile flowers, with sepals and carpel-like organs but no stamen, develop near the apex. The run of these abnormal flowers is followed by an increase in severity of the shoot-like flowers until the end of growth. The *lfy-26 dcl1-7* double mutant plants have approximately as many vegetative leaves, and are as late flowering as *dcl1-7* single mutants (Fig. 3D).

**Figure 3.**
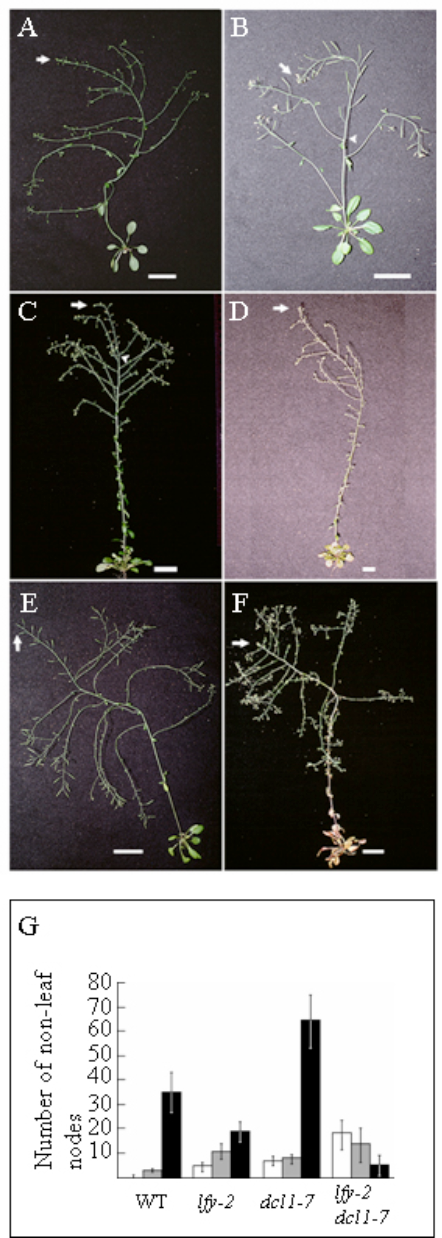
Interaction of *dcl1* with *lfy*. A. *lfy-26*; B. Wild type Co-O; C. *dcl1-7*; D. *lfy-26 dcl1-7* double mutant; E. *lfy-2*; F. *lfy-2 dcl1-7* double mutant. All size bars represent 2.5 cm. Arrows denote the main inflorescence apex; arrowheads, the last lateral coflorescence (in B and C). G. Enhancement of a weak *lfy-2* phenotype by *dcl1-7*. Number of shoot-like organs extending less than 2.5 cm are represented by the white bars; the number of shoot-like lateral branches extending over 2.5 cm are represented by the gray bars; and the number of solitary flowers are indicated by the black bars. Results were collected from sixteen plants in each phenotypic class, except for wild type (+), for which results were collect from ten plants. Double mutants show a tendency to transition away from floral fates towards inflorescence fates, as seen by the increase in the number of shoot like lateral branches, and decrease in the number of solitary flowers.

Lateral organs produced past the vegetative rosette leaves in *lfy-26 dcl1-7* are similar to those of *lfy-26* mutants; however, in the double mutant there is an increased tendency of the lateral primordia to assume a coflorescence fate over the floral fate. This subtle enhancement of the Lfy^-^ phenotype is noticeable as an increase in the number of coflorescence-like branches, each subtended by a cauline leaf, and a reduction in the number of Lfy^-^ flowers along the primary inflorescence axis near the apex.

The weak *lfy-2* mutation converts the first several ‘flowers’ to a more coflorescence-like fate (Fig. 3E) producing 11 ± 2 (*n* = 11) lateral branches subtended by cauline leaves. A wild type plant produces 4 ± 0.5 (*n* = 10) lateral branches before making single flowers. This effect decreases acropetally, and unlike in *lfy-26*, many normal flowers are formed. By contrast, *dcl1-7 lfy-2* double mutants retain the extra leaves and late bolting phenotype of Dcl1^-^ plants. However, during the reproductive phase, the conversion of lateral primordia in *lfy-2 dcl1-7* double mutants from a floral fate to the coflorescence-like fate was extreme, producing a phenotype where nearly all flowers are changed to shoot-like organs (Fig. 3F). The lengths of the lateral organs were classified into three categories (Figure 3,G): shoot-like lateral organs (less than 2.5 cm long); lateral coflorescences (over 2.5 cm); and solitary flowers. Lfy^-^ and Dcl1^-^ plants produce similar numbers of inflorescence shoots and shoot-like lateral organs. In *lfy-2 dcl1-7* segregants, however, the conversion of lateral organs to a coflorescence-like fate was significantly higher than in either single mutant. This can be seen as a sharp decrease in the number of single flowers in the double mutants (Fig. 3G). The enhancement of *LFY* phenotype by *dcl1* is consistent with the notion that *DCL1* controls *LFY* activity either directly or indirectly and by doing so it affects lateral meristem fate transition from coflorescence to flower.

### Early Flowering by *LFY* Over-expression is Enhanced by *dcl1* Mutation

Expression of *LFY* cDNA from the 35S promoter (*35S::LFY*) causes early flowering [37]. If *DCL1* controls *LFY* activity by producing guide miRNA against a repressor of either *LFY* or of its downstream target, then *dcl1-7* mutants harboring *35S::LFY* should flower less readily than *DCL1 35S::LFY* plants. To test this prediction we constructed plants heterozygous for *dcl1* and hemizygous for *35S::LFY* and analyzed the progeny of these self-crossed plants. Table I illustrates the segregation of phenotypes in one representative F_2_ population.

**Table I.** Synergistic interaction between *LFY* gain-of-function and *dcl1* loss-of-function mutations. F2 populations were scored for average flowering time and mean leaf number. The data are presented as mean ± SD.

| Cross | Phenotype | N | *DCL1*<br>Genotype | *35S::LFY*<br>present? | Average<br>flowering<br>time<br>(days) | Average<br>leaf<br>number |
| --- | --- | --- | --- | --- | --- | --- |
| *35S::LFY* x<br>*dcl1-7* | Early<br>flowering | 102 | +/? | + | 21.5 ±2.5 | 7.6 ±1.1 |
|  | Wild type | 22 | +/? | - | 23.7 ±3.9 | 8.5 ±1.1 |
|  | Novel | 24 | *dcl1-7/dcl1-7* | + | 16.4 ±2.6 | 3.8 ±1.0 |
|  | Dcl1<sup>-</sup> | 14 | *dcl1-7/dcl1-7* | - | 37.8 ±9.2 | 17.8 ±3.8 |
| *35S::LFY* x<br>*dcl1-8* | Early<br>flowering | 56 | +/? | + | 22.8 ±3.1 | 7.3 ±1.3 |
|  | Wild type | 17 | +/? | - | 22.0 ±2.5 | 7.6 ±1.3 |
|  | Novel | 14 | *dcl1-8/dcl1-8* | + | 23.1 ±4.0 | 6.9 ±2.3 |
|  | Dcl1<sup>-</sup> | 6 | *dcl1-8/dcl1-8* | - | 34.6 ±4.2 | 16.4 ±3.0 |

Four phenotypic classes were identified: typical *35S::LFY*, wild type, an unexpected novel phenotypic class, and typical Dcl1^-^. The segregation data show no deviation from the 9:3:3:1 ratio for two independently segregating loci (χ^2^ = 2.4; *P* ≥ 0.5), implying that the novel phenotypic class is due to the combined effects of the transgene and *dcl1* mutation. The novel phenotype is characterized by an extremely rapid conversion of the shoot apex to the floral fate, following only three vegetative leaves (Fig. 4A, B; Table I), with no extension of a primary inflorescence stem. A mass of incomplete flowers develops at the rosette level, many with abnormal organs of mosaic tissues. The sepals and petals are often strongly carpelloid (bearing abnormal ovules typical of Dcl1^-^ phenotype), stamens are reduced in size and number, and the carpels are small and often unfused. Similar results were obtained in crosses of the less severe *dcl1-8* allele to *35S::LFY*, with a corresponding weakening of the synergistic phenotype (Table I).

**Figure 4.**
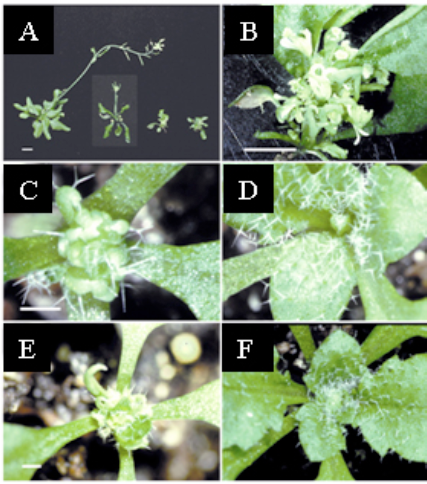
Expression of *LFY* cDNA in a *dcl1* background causes a novel phenotype. A, From left to right, a weak *35S::LFY*, a strong *35S::LFY*, and two examples of *35S::LFY dcl1-7* plants. B, C, and E, Higher magnification of the same *35S::LFY dcl1-7* plant as shown third from the left in panel A: B, at 37 days; C, at 14 days; and E, at 19 days. (D, F) Apex of the weak *35S::LFY* plant shown on the extreme left in panel A: D, on day 14; and F, on day 19. Size bars: A and B = 1 cm; C and D = 1.5 mm; E and F = 5 mm.

To further analyze the synergistic phenotype, individual *35S::LFY dcl1-7* plants were examined daily. By 14 days after germination, the shoot apex of *35S::LFY dcl1-7* terminally commits to the floral fate (Fig. 4C). *DCL1 35S::LFY* plants produce only vegetative leaves at this stage (Fig. 4D). By 19 days, the *35S::LFY dcl1-7* plants produce several carpels, some of which are unfused. Surrounding the carpels is an area that is generally densely packed with many abnormal organs, with combined leaf and floral characteristics or only floral characteristics (Fig. 4E). By this same time, the SAM of most *35S::LFY* plants has finished producing leaves (Fig. 4F) and stem extension has begun. Rosette flowers, typical of *LFY* over-expression, are often seen in the axils of the vegetative leaves by this time. Some variability in the extension of the primary inflorescence was noticed, but no *35S::LFY* plant was ever as severely affected as the *35S::LFY dcl1-7* plants.

To determine the morphological basis of the novel phenotype, the shoot apex was examined on various days post-germination (Fig. 5). Termination of the primary SAM can be seen in *35S::LFY dcl1-7* plants as early as 11 days (Fig. 5A). Confocal fluorescence microscopy of inflorescence stained with propidium iodide revealed the presence of many apical meristem domes instead of a single dome of the SAM present in wild type or *dcl1* mutant plants (Fig. 5G, H). In *dcl1-7 35S::LFY* plants, however, each of the numerous apical domes is smaller in size than in the wild type.

**Figure 5.**
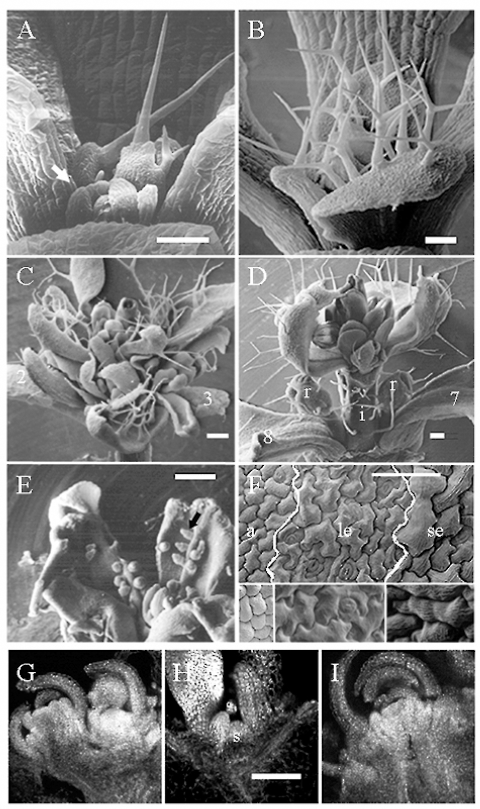
Over proliferation of the SAM in *35S::LFY dcl1* plants. A, C, and E, scanning EM of *35S::LFY dcl1-7* shoot apices: A, after growth for 11 days; C, 16 days; and E, 30 days. Note Dcl1^-^ ovules (filled arrow), with unexpanded integuments in opened carpels in panel E. B and D, shoot apices of *35S::LFY* plants: B, following growth for 11 days and D, 16 days. F, shows the surface features of a single mosaic organ in a *35S::LFY dcl1-7* flower. White lines demarcate the three distinct surface zones that correspond (from left to right) to typical anther, leaf, and sepal, respectively. Corresponding wild type surfaces are shown in the three insets. G-I, confocal laser-scanning images of propidium iodide stained apical meristem: G, *35S::LFY dcl1-7* on day 11; H, wild type on day 11; and I, *35S::LFY* on day 13. Abbreviations: a, typical anther surface; i, inflorescence; le, typical leaf surface; r, rosette flower; s, SAM, and se, typical sepal surface. Numbers indicate the leaf number. Size bars: in A and B, 100 μm; C and D, 200 μm; E, 500 μm; F, 50 μm; G-I, 100 μm.

The fate of the shoot apex does not change from day 11 to day 13 in *35S::LFY* plants, because, at both time points, they produce only leaf primordia (compare 5B with 5I). By 16 days post-germination, the SAM of the *35S::LFY dcl1-7* is clearly consumed in the production of masses of floral organs (Fig. 5C). Some of these floral organs appear normal but misplaced, while others are chimeras of leaf, sepal and sometimes filament cells (Fig. 5F). By this time, *35S::LFY* plants usually show extension of a primary inflorescence that terminates into flowers (Fig. 5D). After approximately 30 days of growth, these *35S::LFY* flowers often contain abnormal organs that have mixed leaf/sepal characteristics and unfused carpels. Similarly, dissected apical regions of 30-day-old *35S::LFY dcl1-7* plants show abnormal ovules typical of Dcl1^-^ plants in malformed carpels (Fig. 5E).

It is possible that the effects of *dcl1* on *35S::LFY* are due to *dcl1* directly affecting *LFY* mRNA. To test this, gel blots of RNA isolated from leaves of these plants were probed for *LFY* mRNA (Figure 1, C). Vegetative leaves of wild type plants express low levels of *LFY* mRNA that cannot be detected by RNA gel blot experiments [38,39]. We found no consistent reduction of *LFY* mRNA in *35S::LFY dcl1-7* plants compared with those in *35S::LFY* plants (Fig. 1, C). One specific *35S::LFY* line did show a high expression of *LFY* mRNA, but it had no phenotypic difference compared the other *35S::LFY* lines. These results do not reveal a consistent direct effect of *dcl1* mutation on the transcript level off *LFY* cDNA.

To examine whether *DCL1* is important for flower initiation via a parallel pathway involving the *EMBRYONIC FLOWER1* (*EMF1*) and *EMBRYONIC FLOWER2* (*EMF2*) genes [40-44], we constructed *dcl1-7 emf1-1*, *dcl1-7 emf2-1*, *dcl1-8 emf1-1*, and *dcl1-8 emf2-1* double mutant combinations. The double mutant phenotypes were indistinguishable from *emf1-1* or *emf2-1* single mutant phenotypes, respectively, suggesting that *DCL1* does not participate in the pathway controlled by *EMF* genes (data not shown).

## Discussion

The control of programmed transition of meristem fate in flowering plants has been an area of intensive study, and a complex pathway that integrates multiple environmental and developmental signals has been elucidated [45–57]. In summary, environmental inputs, including light and temperature, are transduced through a network of interacting genetic pathways to the ‘floral pathway integrator’ genes that include *AP1* and *LFY* transcription factors. These transcription factors in turn activate the genes that initiate reproductive development.

### *DCL1* Augments Flowering through a Pathway Involving *LFY*

We had previously shown that *dcl1-7* and *dcl1-8* mutants, bearing missense mutations in the DExH box RNA helicase domain [7], have delayed onset of the reproductive phase in both long and short days, and cannot be rescued by either vernalization or repeated additions of gibberellic acid [29]. Here we have shown that DCL1 protein is localized in the L2 and L3 layers of inflorescence and floral meristem, and is also expressed in the SAM, ovule integuments, in early but not late embryos, and in the ovule funiculus. We have presented genetic evidence consistent with the idea that *DCL1* participates in the transition of inflorescence to floral fate, presumably along a pathway that involves *LFY*. *DCL1* does not appear to have a direct genetic interaction with *LFY*, nor does it directly regulate *LFY* mRNA.

Results of double mutant analysis are most consistent with *DCL1* being a negative regulator of a putative repressor of *LFY*, probably by catalyzing the formation of a guide miRNA against the repressor. However, this role alone does not explain the synergistic phenotype exhibited by *dcl1* mutant plants that ectopically expressed *LFY*. To reconcile these results we propose that the synergistic effect is due to the secondary effect of *dcl1* mutation on cell proliferation in the SAM. Genetic analysis had previously indicated that *DCL1* controls the extent of cell division in the SAM [6]: *dcl1* mutants have increased proliferation of meristematic cells. We propose that in a *dcl1* mutant harboring the *35S::LFY* gene, the early exposure of more numerous SAM cells to the adventitious expression of *LFY* mRNA recruits additional cells into the floral fate, producing numerous early flower or flower-like organs at the apex. Consistent with *DCL1*’s proposed effect on meristem cell proliferation is our finding that DCL1 protein is localized on the SAM. Expression of *DCL1* cDNA, however, did not increase the amount of protein in the SAM, which may be due to feedback regulation of *DCL1* mRNA through miR162 [25,32], or it may reflect a combination of transcriptional and miRNA-mediated post-transcriptional regulation.

### *DCL1* in Early Embryo

Mutations in *dcl1* show sporophytic maternal inheritance [28], in which even a heterozygous embryo borne on a homozygous mutant flower is defective in pattern formation, thus leaving open the possibility that the observed maternal effect could be due to the transmission of DCL1-generated miRNA through the female gametophyte into the egg. However, a *DCL1* promoter fragment of the maternally transmitted copy (but not the paternal copy) is transcriptionally active in the embryo [7], suggesting that at least in principle some DCL1 transcripts in the embryo could be synthesized off the maternally inherited allele. Our ability to detect the presence of DCL1 protein in early embryo cells provides evidence that in principle some miRNA could be synthesized in these cells *de novo* by the action of DCL1 protein on miRNA precursor molecules. These results are consistent with the embryo-lethal phenotype of homozygous *dcl1* deletion mutants [3] in the sense that DCL1 protein in early embryo is essential for viability, presumably because it is necessary for *de novo* miRNA synthesis. This does not eliminate the possibility that at least some mature miRNA, or even all pri-miRNA/pre-miRNA precursors, in the early embryo are maternally transmitted.

## Materials and Methods

### RNA Analysis

Methods for RT-PCR analysis to detect the 6.2 kb *DCL1* transcript were previously described [7], except that radio-labeled nucleotides were incorporated at the PCR step. Primers for amplifying a 465-bp fragment of *DCL1* were 5’-d[GGGTCAATGGTGTGCTTACAAGG]-3’ and 5’-d[CACCGTATAAAACTAAGCGAAGGCAGC]-3’; for a 523-bp fragment of β-1 tubulin gene were 5’-d[CGTAAGGAAGCTGAGAACTGTGATTGCC]-3’ and 5’-d[CGTCCCACATTTGCTGTGTCAGC]-3’; and for a 746-bp fragment of *ROC1* gene were, 5’-d[CCTCTTCTTCAGTCTGATAGAGATC]-3’ and 5’-d[GAGTGCTCATTCCTTATTTCTGG]-3’. After 20 amplification cycles, the samples were fractionated by electrophoreses through a TBE polyacrylamide gel, which was subsequently dried down, and phosophorimaged. No signal was detected in samples without reverse transcriptase. For RNA gel blot analysis, the probe for *LFY* template was a 900 bp HinD III*-* BamH I fragment of pDW124 [37], which hybridizes to a *LFY* mRNA of 1.5 kb, and for *ROC1* template was a 400 bp PCR fragment from pCG22 (courtesy Dr. C. Gasser), amplified with primers 5’-d[GATCGTGATGGAGCTGTAC]-3’ and 5’-d[CAATCGGCAACAACCAC]-3’, which hybridizes to a constitutively expressed 750 bp cyclophilin mRNA [48].

### Protein analysis

To generate antibody against the unique N-terminal fragment of DCL1 protein, the corresponding cDNA sequence region was amplified with primers 5’-d[TTATATCTAGACATGGTAATGGAGGATGAGCCTAGA]-3’ and 5’-d[TTCGTCAAGCTTACCGCCATTCTTTTGCAACCCATT]-3’, cloned into pGEX-KG (*Xba*I-*Hin*dIII) in *E. coli* BL21, the resulting in-frame fusion with glutathione S-reductase (GST) epitope was confirmed by complete sequencing, the fusion protein (37 kd) expressed and purified by affinity chromatography on glutathione-sepharose, confirmed by SDS-PAGE and Western blot analysis. Approximately 6 mg purified protein was concentrated, and rabbit antibodies were raised by Alpha Diagnostics (San Antonio, Texas). Antigen-affinity purified antibody was obtained from the polyclonal antiserum as follows. The N-terminal DCL1 fragment expressed as a GST-fusion protein (15 mg) was coupled (98% efficiency) to Aminolink Immobilization affinity column (Pierce), 1 ml of polyclonal antiserum was applied to the column, washed successively with low and high salt buffers. The bound antibody was eluted with 10 ml of 100 mM triethylamine (pH11.5); eleven 1 ml fractions were immediately neutralized and analyzed by Western blots following SDS-PAGE. The sixth fraction eluted from the column contained pure anti-DCL1 antibody (∼100 *µ*g/ml, data not shown), which was used in all immunolocalization experiments. As a negative control, 0.5 ml of preimmune antiserum diluted with 9.5 ml ImmunoPure IgG binding buffer (Pierce) was purified on Protein A/G affinity column (Pierce) according to manufacturer’s instructions. The bound antibody was eluted with 5 ml of ImmunoPure® IgG elution buffer; of the six 1 ml fractions collected, the sixth fraction was free of sera proteins (∼100 *µ*g/ml, data not shown) and was used for all immunolocalization experiments. Leaves and buds of Arabidopsis (Co-O) were collected from plants that had just started flowering, 25 g of tissues were ground in liquid nitrogen, extracted with 40 mM Tris, pH 9.5; 50 mM MgCl_2_, 2% PVPP, 1 mM PMSF, and EDTA-free complete protease inhibitor (Roche), centrifuged at 12,000x g for 10 min at 4°C. The supernatant was precipitated with an equal volume of ice-cold acetone and incubated at –20°C for 2 h. The precipitated protein was centrifuged for 15 min at 12,000 g, the pellet washed with 70% acetone, air-dried and dissolved in 500 *µ*l of 50 mM Tris-HCl buffer (pH 7.5) containing 150 mM NaCl and protease inhibitors. The protein was immunoprecipitated with DCL1 specific antibody. 1 μg of antibody was added to 500 *µ*l of acetone-precipitated protein and incubated at 4°C overnight in an end-to-end mixer. The immuno-complex was pulled down by adding 20 *µ*l of Protein G sepharose beads (Amersham Biosciences) for 1 hr at 4°C. The beads were collected by centrifugation at 2,500 rpm for 5 minutes at 4°C, followed by 4 times wash in 10 mM Tris-HCl (pH 8.0) containing 250 mM LiCl, 0.5% (w/v) Sodium deoxycholate and 1 mM EDTA. The washed beads were boiled in 50 *µ*l SDS sample buffer and separated on 4-20% (w/v, acrylamide) SDS-PAGE gradient gel. The separated proteins were transferred to polyvinylidene fluoride membrane (PVDF, Bio-Rad) and probed with rabbit anti-DCL1 antiserum, or with rabbit pre immune serum as a negative control. Western blot analysis was performed according BioRAD, antibody complexes were visualized with either alkaline phosphatase conjugated substrate kit (BioRad) in a colorimetric reaction or with SuperSignal West Dura Substrate Solution (Pierce) and exposed to X-ray films. For immunocytochemistry, plant tissues were fixed, embedded, sectioned and de-waxed as described by [7]. The sections were rehydrated, blocked in 3% (w/v) bovine serum albumin (BSA), incubated in 1:100 dilution of the primary affinity purified antibody in 3% BSA overnight, washed five times in phosphate buffered saline, incubated with 1:100 dilution of secondary antibody overnight, and washed as before. The slides were developed using the Vectastain ABC and Vector VIP kits (Vector Laboratories, Inc.) according to manufacturer’s instructions.

### Strains and Genetic Methods

Strain Sin1-C (*dcl1-7/+*; *gl-1*) was previously described in [31]. For constructing *lfy-26 dcl1-7* double mutants, pollen from Sin1-C was crossed to CS6296 (*lfy-26*/*+*; *ap1-1*; *er*; Landsberg ecotype, *La-O*). F_1_ plants were distinguished from selfed progeny by their Ap1^+^ phenotype. F_2_ lines segregating Lfy^-^ plants were chosen for analysis. Double mutant *ap1 dcl1* plants were identified by their enhanced phenotypic effects [29]. Phenotypic analysis was done exclusively on *ERECTA* plants. To make *lfy-2 dcl1-7* double mutants, pollen from Sin1-C was crossed to CS6229 (*lfy-2*; Columbia ecotype, *Co-O*). For quantitative estimation of phenotypes in *lfy-2* x *dcl1-7* crosses, lateral organ lengths were measured during growth of plants. Status of the *dcl1-7* allele was checked by linkage to nga59, a marker that shows 100% linkage with *dcl1* [7]. Pollen from *dcl1-7* was crossed to DW151.202, a strain containing the *35S::LFY* transgene [37]. The synergistically affected plants are inferred to be *dcl1* because they were homozygous for the closely linked *La-O* allele of the nga59 SSLP marker, contained Dcl1^-^ ovules, and segregated at the expected frequency. Presence of the *35S::LFY* transgene was verified by PCR assays. Control crosses of *dcl1-7* to the *No-O* background indicated that there is no suppressor of *DCL1* present in *No-O* (see below). To make *dcl1 emf* double mutants, pollen from *dcl1-7* was crossed to plants heterozygous for *emf1-1* and *emf2-1* (Co-O) [40-42]. The F2 progeny in crosses segregating *emf* were tested for the *dcl1* genotype by association with nga59 PCR. Conditions for plant germination and growth were as described before [29,31]. Transgenic lines expressing *DCL1* full length cDNA under cauliflower mosaic virus 35S RNA promoter and their construction were described earlier [32].

### Microscopy

Techniques for light and scanning electron microscopy were as described [29,31]. Confocal microscopy was performed as described [49] using a Leica DM-IRB microscope with 40x-oil immersion lens on tissues stained with propidium iodide. Single plane optical sections were obtained (60 MW Argon laser at 488 nm, LP550 filter, and set to slow scan) at 2 - 4 accumulations per section.

## Acknowledgements

We thank Detlef Weigel, Martin Yanofsky, CharlesGasser, and Alan Lloyd for strains and plasmids; Arabidopsis Stock Center for strains; Terry Delaney and Lynne Angerer for advice and comments; Marybeth Langer for discussion; Sarah Bean, Jack Tsai, Joanne Lundholm, and Patricia Gilligan for assistance.

## Authors’ contributions

SES performed RNA analysis, antibody purification, immunocytochemistry, confocal microscopy, and wrote part the first draft; TAG performed strain construction, performed genetic analysis, and helped writing the first draft; BNP re-purified the antibody and performed protein analysis on plants; JDL did strain constructions and performed genetic and phenotypic analysis, SR performed scanning E/M; DSM assisted SES in constructing and expressing DCL1 fragments for antibody production as well as antibody purification and immunocytochemistry; DC and BC advised SES on antibody purification; AR planned and supervised the project, participated in genetic and phenotypic analysis, procured funding, wrote and edited the manuscript.

## Funding

This work was supported by NSF grants IBN-9728239, IBN-9982414, EIA-0130059, EIA-0205061, and FIBR-0527023 to A.R.

